# Maternal transfer of mRNA LNP-derived, pathogen-specific, monoclonal IgG to suckling mice

**DOI:** 10.1101/2025.03.12.642889

**Authors:** Jennifer E. Doering, Yetunde Adewunmi, Cailin E. Deal, Obadiah Plante, Andrea Carfi, Nicholas J. Mantis

## Abstract

Breast milk provides a rich source of naturally derived maternal antibodies that confer passive immunity to infants, protecting them from a variety of respiratory and enteric infections. For at-risk newborns in low- and middle-income countries, supplementing breast milk with pathogen-specific neutralizing and bactericidal antibodies could offer significant short- and long-term health benefits. In this study, we explored the use of mRNA and lipid nanoparticle (LNP) technology to deliver a Vibrio cholerae-specific monoclonal IgG antibody (“ZAC-3”) into the milk of lactating mice. Swiss Webster mice were intravenously administered ZAC-3 IgG mRNA-LNPs, and we monitored serum and breast milk for the presence of V. cholerae-specific human IgG1. A single injection of mRNA-LNPs led to rapid and sustained expression of ZAC-3 IgG in both the blood and breast milk of lactating dams. ZAC-3 IgG1 in these samples recognized whole V. cholerae cells by ELISA and exhibited potent vibriocidal activity in the presence of human complement. Furthermore, ZAC-3 IgG was detected in the sera of suckling pups at levels proportional to those in the mothers, demonstrating successful transfer of functional antibodies to the newborns. In conclusion, our findings highlight the potential of mRNA-based monoclonal antibody platforms in the maternal-newborn context and address key challenges associated with the direct delivery of recombinant antibodies.

## INTRODUCTION

Worldwide, children under the age of five remain at risk of infectious diseases affecting the lower respiratory tract, bloodstream (neonatal sepsis), or gastrointestinal system (diarrhea) that can have long term consequences for health and development. Breastmilk provides a crucial source of passively immunity during the early days, weeks and even months of life, as it contains an array of antibodies against viral and bacterial pathogens that the mother has encountered through infection or active vaccination [1]. For example, a recent examination of breastmilk samples collected from women in 5 high and low-to-middle income countries revealed IgA and IgG antibodies against hundreds of pathogen-derived antigens [2]. The observed profiles are consistent with observations dating back decades that breastfed infants have reduced risk of infections, hospitalizations and morbidity as compared to non-breastfed cohorts [3]. Breast feeding also influences the gut microbiota and has health benefits on immune development [4]. Proactively modulating specific antibody titers in colostrum and breastmilk in anticipation of potential pathogen exposures represents a powerful means to improve newborn health.

Augmenting pathogen-specific antibody levels in serum as well as colostrum and breastmilk can be achieved by immunizing women during pregnancy [5]. For instance, mothers who received influenza vaccines in the third trimester of their pregnancy maintained high levels of IgA in their breastmilk [6]. Likewise, abundant levels of pertussis-specific IgG and IgA are detected in breastmilk after maternal vaccination during pregnancy. [7; 8]. However, in some cases, the concentration of maternal antibodies passed to the offspring is insufficient or declines too quickly to afford protection [9]. Moreover, in the case of a rapidly spreading disease outbreak or in instances where effective vaccines do not exist, an alternative strategy of providing immunity to newborns is required. In such situations, monoclonal antibodies (mAbs) could provide immediate protective immunity.

There are pharmacokinetic and clinical pharmacology challenges associated with the use of mAbs directly in children such as dosing complexities, bio-distribution, route of administration and immunogenicity [10]. Currently, only a few mAbs such as Palivizumab and Nirsevimab, which neutralize respiratory syncytial virus (RSV), are approved by the FDA for direct use in neonates [11]. This is because, generally, the use of mAbs in pediatrics is only explored after their safety and efficacy has been well established in adults. Given that maternal mAbs in breastmilk can also attach to, and swiftly eliminate, pathogens found in breastfeeding infants, using breastmilk as a vehicle to deliver pathogen-specific monoclonal antibodies could provide a means to safely boost antibody levels by mimicking natural passive immunity [12]. The expression of mAbs from mRNA has demonstrated potential in animal models for diseases such as chikungunya, HIV-1, and SARS CoV-2, with protective levels observed in mice, non-human primates and humans [13]. In this study, we implemented this therapeutic approach to directly deliver functional antibodies to infants through breastmilk, presenting it as a promising method to restrict pathogen replication and safeguard newborns from diseases. We employed cholera as a proof of concept, as it is a disease characterized by recurrent outbreaks and disproportionally affects children who would benefit greatly from a passive immunotherapy strategy. ZAC-3 is a well-established protective mAb that targets the conserved core region of lipopolysaccharide (LPS) in *Vibrio cholerae* O1 strains. Our objective was to evaluate whether mRNA-encoded ZAC-3 IgG1, administered to mice, could be expressed and subsequently transferred to their offspring through breastmilk. Here, we demonstrate that systemic immunization with mRNA/LNP resulted in robust expression of functional ZAC-3 IgG in the serum of adult mice, a portion of which was successfully transferred to breastmilk and serum of suckling mice.

## MATERIALS and METHODS

### Recombinant antibodies

Recombinant ZAC-3 IgG was produced by GeneArt (Thermo Fisher Scientific, Waltham, MA). ZAC-3 IgG was purified from transient transfection of a 2:1 ratio of HC to LC by weight in EXPI293 with HiTrap Protein A and polished using HiLoad Superdex 200 26/600 prep. ZAC-3 IgG concentrations were determined by absorption at 280nm using a NanoDrop microvolume spectrophotometer (Thermo Fisher Scientific).

### Generation of modified mRNA and LNPs

Sequence-optimized mRNA encoding functional ZAC-3 IgG was synthesized *in-vitro* using an optimized T7 RNA polymerase-mediated transcription reaction with complete replacement of uridine by N1-methyl-pseudouridine and a DNA template with open reading frames flanked by 5’ untranslated region (UTR) and 3’ UTR sequences with a terminal encoded polyA tail [14]. Lipid nanoparticle-formulated mRNA was produced through a modified ethanol-drop nanoprecipitation process as described [15]. Briefly, ionizable, structural, helper and polyethylene glycol lipids were mixed with mRNA (2:1 HC to LC for IgG) in acetate buffer at a pH of 5.0 and at a ratio of 3:1 (lipids:mRNA). The mixture was neutralized with Tris-Cl at a pH 7.5, sucrose was added as a cryoprotectant, and the final solution was sterile filtered. Vials were filled with formulated LNP and stored at −70°C. The drug product underwent analytical characterization, which included the determination of particle size and polydispersity, encapsulation, mRNA purity, osmolality, pH, endotoxin and bioburden, and deemed acceptable for *in-vivo* study.

### Bacterial strains and growth conditions

*V. cholerae* 01 strains O395 (classical Ogawa) and C6706 (El Tor Inaba) are described in **Table S1**. Bacteria were routinely cultured in LB broth at 37°C with aeration. As necessary, media were supplemented with streptomycin (100 μg/ml).

### Mouse experimental protocols

Nonpregnant and timed pregnant female Swiss Webster (SW) mice aged 6 to 12 weeks were obtained from Taconic Biosciences (Germantown, NY). All experiments were performed in accordance with protocols approved by the Wadsworth Center’s Institutional Animal Care and Use Committee. LNPs (1 mg/kg) or sterile PBS were administered by intravenous (IV) tail vein injection unless noted otherwise.

ZAC-3 IgG expression was assessed in serum by collecting submandibular blood and spinning samples at 13,000 x g for 10 min before transferring serum to a clean tube. To identify antibody expression in breastmilk, dams were separated from pups 2-4 h prior to sample collection to allow milk to accumulate. At the time of milking, mice were anesthetized with isoflurane and subcutaneously injected with 2 IU of oxytocin. Milk was collected into 15 mL conical tubes using a modified breast pump. Tubes were spun for 20 min at 1,500 x g at 4°C before transferring the whey portion to a fresh tube. A second spin at 13,200 x g at 4°C for 15 min before transferring the aqueous portion to a fresh tube. Total human IgG1 levels in serum and breastmilk were determined using an isotype-specific ELISA, as detailed below.

### Capture enzyme-linked immunosorbent assays (ELISA)

To quantitate human IgG in serum and breastmilk samples, 384 well plates were coated with 0.02 mL per well of goat anti-human IgG Fc fragment (Bethyl Laboratories, Montgomery, TX) at 1:100 dilution in 0.05 M carbonate-bicarbonate buffer and incubated at room temperature for 3 h. The 384 well plates were then washed with 0.05% PBS-T and blocked overnight at 4°C with 0.05 mL per well of 2% goat serum in PBS-T. Serum and breastmilk samples were serially diluted in 2% goat serum in PBS-T and transferred (0.02 mL per well) to the coated plates and incubated at room temperature for 90 min. Plates were then washed and incubated with goat anti-human IgG horseradish peroxidase (HRP; SouthernBiotech, Birmingham, AL; 1:5000) at room temperature for 1 h. Plates were subsequently washed and incubated with SureBlue TMB 1-C substrate (SeraCare, Gaithersburg, MD). The reaction was stopped with 0.02 mL per well of 1 M phosphoric acid and read at an absorbance of 450nm on a SpectraMax iD3 Microplate Reader (Molecular Devices, San Jose, CA). Absolute quantities of human antibody were extrapolated against a standard curve in GraphPad Prism 10.

### Bacterial whole cell ELISA

Immunolon^TM^ 4HBX 96-well microtiter ELISA plates were coated overnight with 200µL of *V. cholerae* C6706 at an OD_600_ of 0.1. The plates were blocked for 2 h with 2% goat serum in PBS-T, washed and then treated with 100 µL serial dilutions of mouse serum samples. After a 1 h incubation at room temperature, the plates were washed and treated with 100 µL of goat anti-mouse HRP secondary antibody (1:2000 dilution; SouthernBiotech,) for 1 h at room temperature. ELISA plates were washed and developed using SureBlue^TM^ Microwell Peroxidase Substrate. Plates were analyzed using a Spectromax 250 spectrophotometer with Softmax Pro 7.0 software (Molecular Devices).

### Complement-dependent serum bactericidal assay

The vibriocidal activity of sera and breastmilk was determined using the agar plate vibriocidal assay. Briefly, an overnight culture of *V. cholerae* C6706 was sub-cultured in LB then grown to mid-log at 37 C with aeration for 2 h. Mid-log phase cells were diluted to 4×10^5^ CFUs/mL in PBS supplemented with guinea pig serum (20% vol/vol) then mixed 1:1 (vol/vol) with mouse milk and serum samples that had previously been incubated at 56 C for 30 min to inactivate endogenous complement in 96-well plates. Plates were then incubated at 37 C for 1 h and 50 µL aliquots were subsequently spread on LB plates. The plates were incubated overnight at 37 C and colonies were counted to determine the residual viability of *V. cholerae* C6706 that had been exposed to serum or milk samples in the presence of guinea pig complement.

### Neonatal intragastric challenge model and protection

Pups were gavaged per os (P.O.) with 0.05 mL of mid-log phase *V. cholerae* C6706 (1 × 10^8^ CFUs/mL) two days (48 h) after their dams had received LNPs or vehicle control. One day (24 h) later, the pups were euthanized by decapitation and blood collected immediately thereafter. The pups were dissected, and the small and large intestines were harvested and placed into 1 mL of sterile PBS, then homogenized (2 x 30 sec) using a Bead Mill 4 Homogenizer (Thermo Fisher Scientific). To enumerate bacterial CFUs in the homogenates, serial dilutions were plated on LB agar supplemented with streptomycin (100 μg/ml) then incubated overnight at 37°C. CFUs were determined manually. From the dams, blood and breastmilk samples were collected on the same day the pups were challenged with *V. cholerae*.

### Statistical analysis

Statistically analysis and graphics were performed using GraphPad Prism version 10.4 (Boston, MA). Differences between groups were determined using an ordinary two-way ANOVA with a Sidak’s multiple comparison test (P<0.05).

## RESULTS

To determine if ZAC-3 IgG1 is successfully expressed from an mRNA template *in-vivo,* we administered nucleoside-modified ZAC-3 IgG1 mRNA LNPs (1 mg/kg) to adult female SW mice intravenously and monitored sera for the onset of human IgG and *V. cholerae*-specific antibodies over seven days (**Figure 1A**). Human IgG expression was detected in all mice at all time points examined, with peak concentrations (>200 µg/mL) at 48 h (**Fig. 1B**). There was no IgG1 detected in sham treated animals (**Fig. 1B**). A bacterial whole cell ELISA confirmed that human IgG1 in serum bound to immobilized *V. cholerae* C6706 (Ogawa), (**Fig. 1C**).

**Figure 1.**
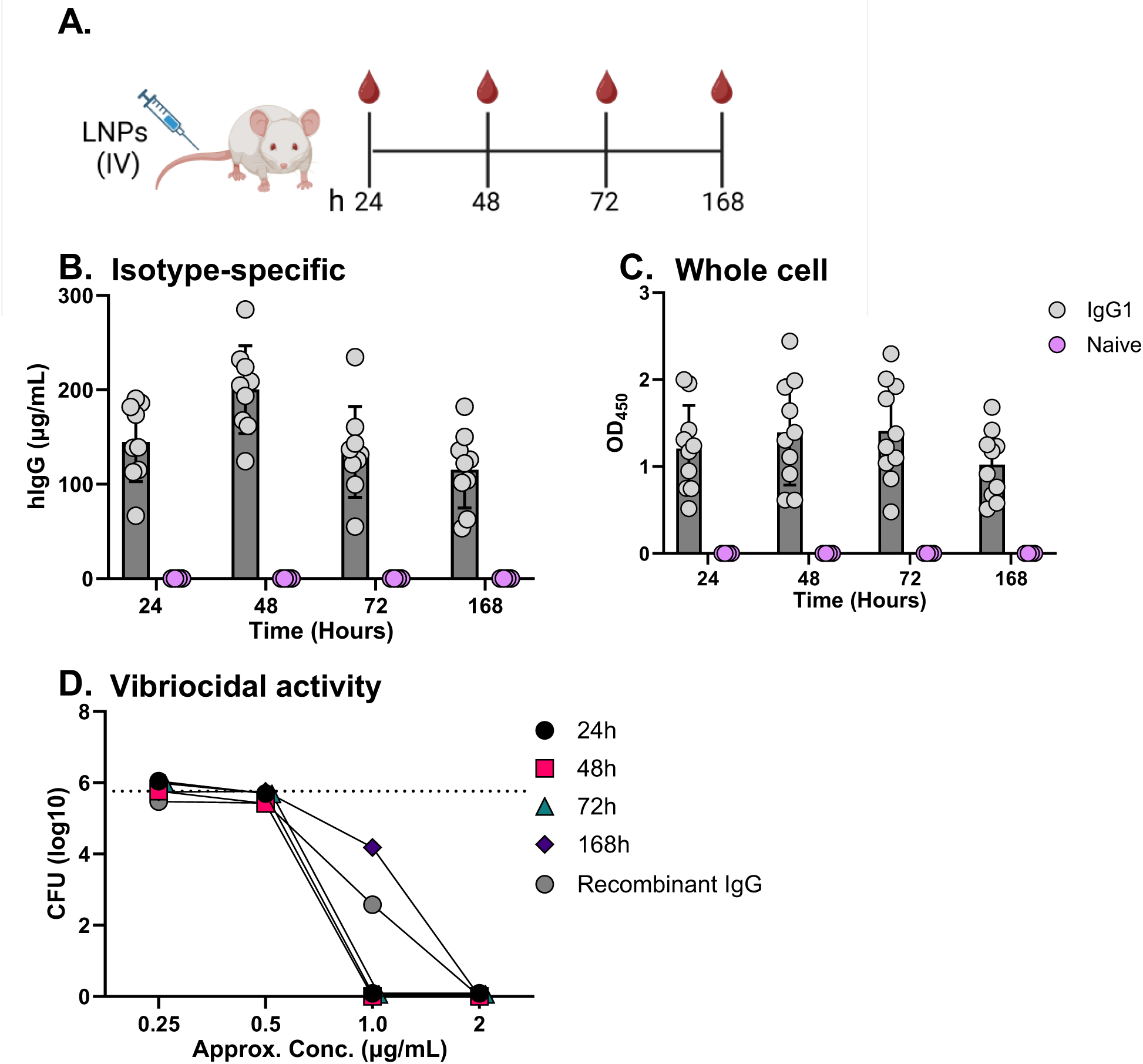
ZAC-3 IgG1mRNA expresses in serum and demonstrates potent vibriocidal activity. (**A**) Adult female Swiss Webster mice were intravenously administered 1 mg/kg of ZAC-3 encoding mRNA-LNPS or PBS and serum was collected at 24, 48, 72, and 168 h. (**B**) Presence of antibody was detected using isotype-specific ELISA. (**C**) *V. cholerae* C6706 specific IgG titers were determined using a whole cell ELISA (serum used a 1:50 dilution). (**D**) Samples within each group were pooled for use in a vibriocidal assay in the presence of guinea pig complement, as described in the Materials and Methods. The dashed line represents the average CFUs recovered from naïve serum as equivalent dilution to experimental samples. The column height represents the mean ± SD per group (n = 4-10 mice per group).

To assess functionality of ZAC-3 IgG1, we performed a modified complement-dependent vibriocidal assay. Importantly, all serum samples containing *V. cholerae-* specific antibodies demonstrated >99.999% vibriocidal activity at 2 µg/mL (**Fig. 1D**). Values were similar to recombinant ZAC-3 IgG1, which elicited vibriocidal activity, while naïve serum resulted in no killing (Fig 1D).

We next examined the presence and functionality of ZAC-3 IgG1 levels in lactating mice and compared levels to non-lactating mice at four different timepoints (24, 48, 72 and 168 h) (**Fig. 2A**). We observed that ZAC-3 IgG1 was five times as high in the sera of non-lactating mice as compared to lactating mice at all time points examined (**Fig. 2B**). The observed differences in ZAC-3 IgG1 concentration were also reflected in the vibriocidal activity of the serum samples, as sera from non-lactating mice was 40% more efficient at killing *V. cholerae* C6706 than sera from lactating dams at equivalent dilutions (**Fig. 2C**). Taken together, these results imply that a large portion of ZAC-3 IgG1 may be shuttled out of circulation during lactation and distributed to other locations such as the mammary glands (**Fig. 2C**).

**Figure 2.**
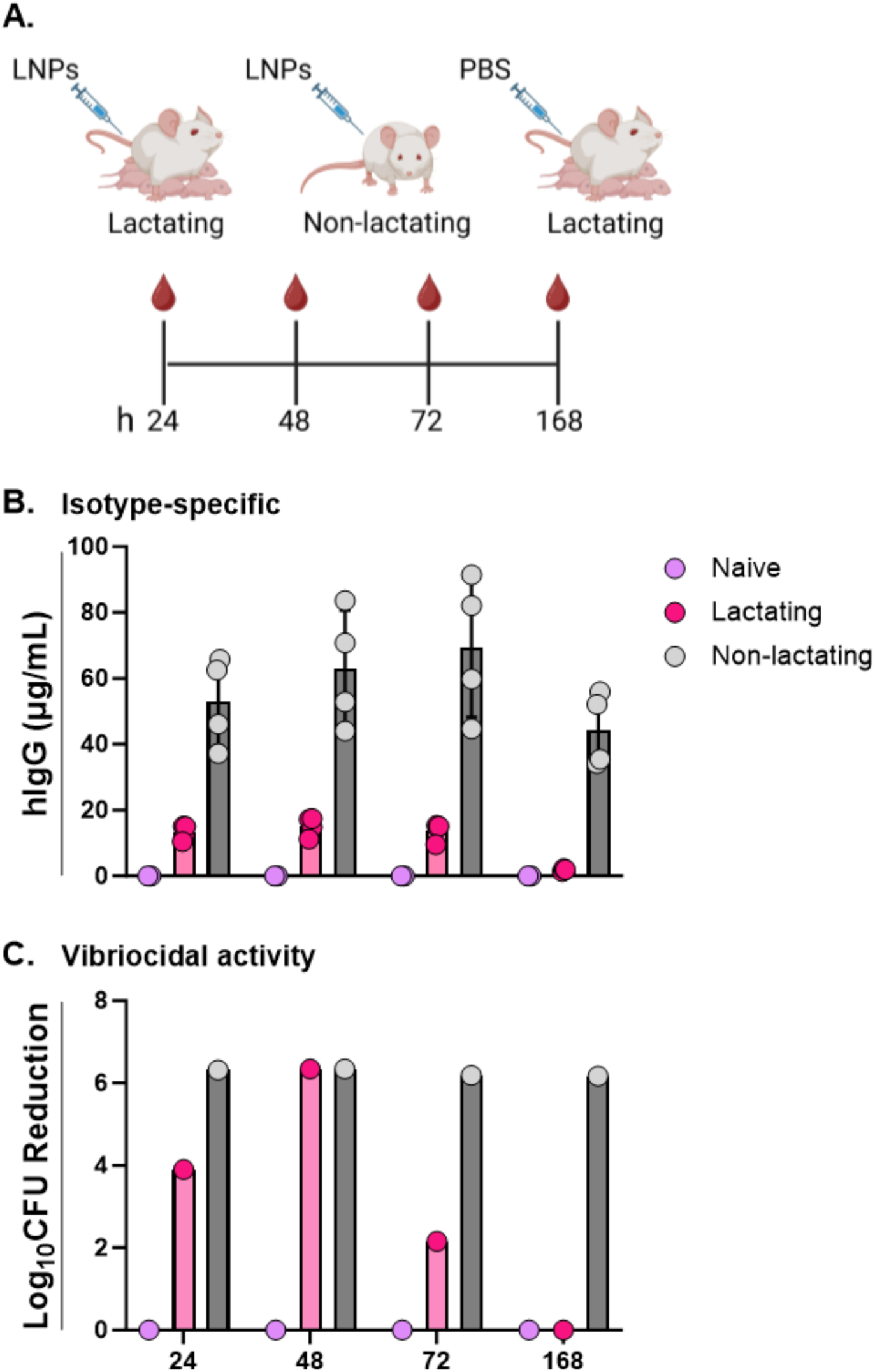
ZAC-3 serum concentrations dependent on lactation status. Lactating and non-lactating Swiss Webster mice were intravenously administered 1 mg/kg of LNPs or PBS 3-7 days after giving birth. (**A**) Cartoon depicting experimental design in which serum was collected at 24, 48, 72, and 168 h. (**B**) Serum hIgG concentrations as interpolated using an isotype-specific ELISA. Each symbol indicates a single mouse. (**C**) Serum samples within each group (naïve, lactating, non-lactacting) were pooled at indicated time points for use in a vibriocidal assay. The column height represents the mean ± SD per group.

One explanation for why ZAC-3 IgG1 levels were lower in the sera of lactating is that a significant portion of IgG1 is being shunted into breastmilk. To test this, we collected milk from lactating dams daily between 24 and 96 h post LNP administration. The whey fraction was analyzed for functional ZAC-3 (**Fig. 3A**). We found that ZAC-3 IgG1 increased progressively over time with an average peak of 2 µg/mL at 96 h post-administration. The estimated amount of ZAC-3 IgG in milk was ∼5 times less than in serum (**Fig. 2B, 3B**). This suggests that a fraction of the ZAC-3 IgG in circulation is directed to the mammary glands. Nonetheless, at a 1:4 dilution, ZAC-3 present in each milk sample was able to bind to immobilized *V. cholerae* C6706 and as expected, the strength of the binding interaction was dependent on the concentration of ZAC-3 IgG1 that was present in the milk samples (**Fig. 3C**). Moreover, pooled milk containing ZAC-3 at 96 h killed 100-fold more *V. cholerae* C6706 than naïve milk samples thus demonstrating that these antibodies were functional and exhibited vibriocidal activity (**Fig. 3D**).

**Figure 3.**
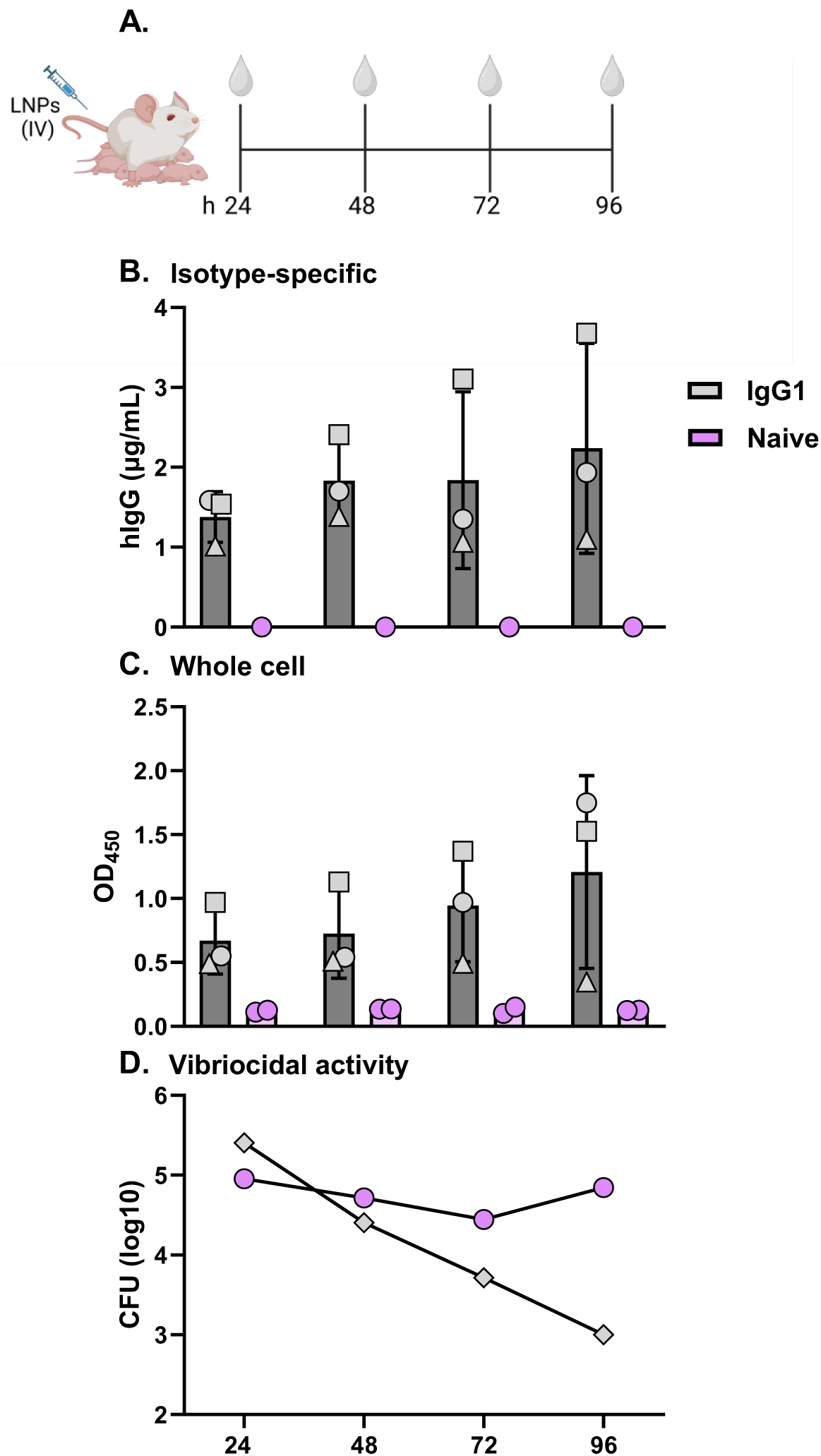
ZAC-3 IgG1mRNA expresses in breastmilk and demonstrated vibriocidal activity. Swiss Webster mice were intravenously administered 1 mg/kg of LNPs or PBS 3-7 days after giving birth. (**A**) Cartoon depicting experimental design in which milk was collected at 24, 48, 72, and 168 h. (**B**) hIgG concentrations detected using isotype-specific ELISA in milk collected at indicated time points from three lactating dams (indicated by circle, square and triangle symbols). (**C**) *V. cholerae* whole cell ELISA (results shown for breastmilk at 1:4 dilution). (**D**) Samples within each group were pooled for use in a vibriocidal assay. Each group represents the mean ± SD of 2-3 mice per group.

Next, we investigated the extent of antibody transfer between dams and pups over the course of 7 days by analyzing serum IgG collected from pups that fed from vaccinated dams (**Fig. 4A**). We tested pup serum as other studies have shown that breastmilk derived antibodies can pass into the serum of breast-fed rodents. As expected, our results show that the concentration of ZAC-3 IgG1 in the serum of pups steadily increased at all time points tested with most plateauing by 168 h with concentrations around 4.5 µg/mL (**Fig 4B**). Although, we observed that the total daily concentration from litter to litter varied (**Fig 4B**), this was not surprising as the concentration of IgG in pup serum was dependent on the available amount of transferrable IgG1 within the breastmilk of each dam (**Fig. 4B**). A single pool of serum from each litter also demonstrated vibriocidal activity against *V. cholerae* C6706 in a concentration-dependent manner with little to no impact at 24 and 48 h but an increasing 3-log fold reduction in CFUs noted from 72 to 168 h (**Fig. 4C**). This confirms that ZAC-3 retains functionality following transfer from dam to pup through ingested breastmilk and vibriocidal activity is dependent upon concentration.

**Figure 4.**
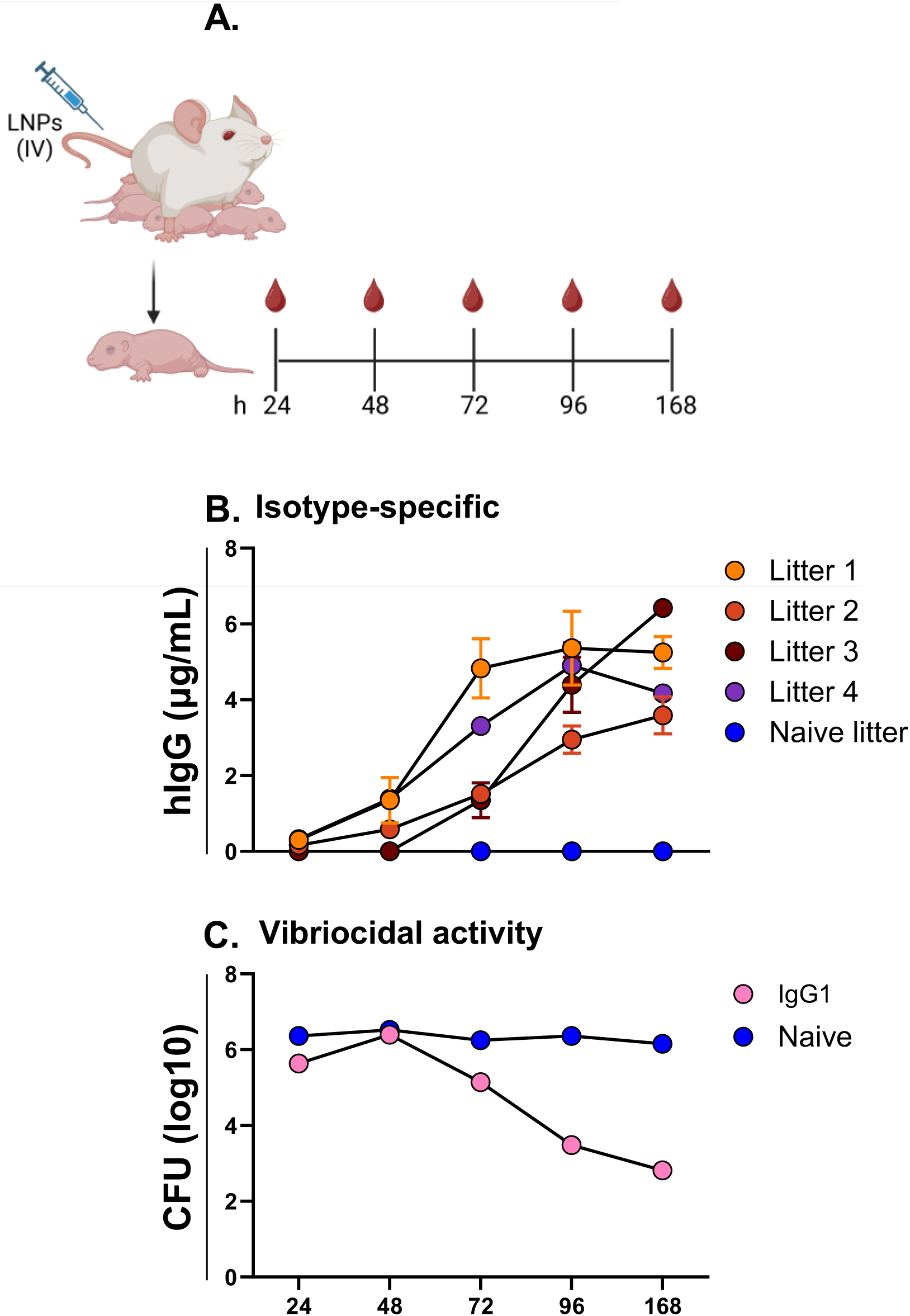
ZAC-3 IgG transferred to neonates accumulates in circulation over time. ZAC-3 mRNA-LNPs (1 mg/kg) or PBS was intravenously administered to dams on 3-7 days after giving birth. This was considered time 0. (**A**) Cartoon depicting experimental design in which serum was collected from pups at 24, 48, 72, and 168 h from four different litters. (**B**) Serum hIgG concentrations as determined using an isotype-specific ELISA. For each time point, a symbol represents 2 pups removed from the litter and euthanized for serum collection. (**C**) Samples from each timepoint (2 mice per litter) and each litter from naïve (n=2) or ZAC-3 mRNA treated (n=4) were pooled and used in the vibriocidal assay, as described previously.

To further probe the functional activity of the ZAC-3 IgG expressed from mRNA in mice, we performed an *in-vivo* neonatal intragastric challenge. Breast-fed pups were intragastrically challenged with 50 µL of 5 X 10^6^ CFUs of *V. cholerae* C6706 48 h after dams were given LNPs or PBS (**Fig. 5A**). Pups were then separated from dams overnight and sacrificed 24 h after challenge. Maternal blood and breastmilk were collected at the time of challenge. Our results show that, consistent with previous experiments, ZAC-3 was expressed in the serum and milk of all dams that received LNPs intravenously (**Fig. 5B**). However, as highlighted earlier, the concentration of ZAC-3 IgG in milk was at least 4 times less than in the serum (**Fig. 5B**). In a similar manner, ZAC-3 IgG was also present in the serum of breast-fed pups at virtually identical concentrations as dam milk, resulting in a 1:1 dam milk to pup serum concentration ratio (**Fig. 5B**). Following intragastric challenge, the log-CFUs of *V. cholerae* C6706 recovered from pup guts did not reveal any significant difference between the pups fed by naïve dams and those fed by dams given LNPs (**Fig. 5C**). Given that antibody concentrations in pup circulation were found to be at or below the threshold for bacterial killing, levels in the gut were most likely too low to provide clearance. ZAC-3 IgG was not detected in gut tissue homogenates derived from suckling pups, possibly because the antibodies were rapidly absorbed into circulation or were below the limit of detection for our binding assay (data not shown). Thus, the *in-vivo* functional activity of ZAC-3 in this model could not be ascertained.

**Figure 5.**
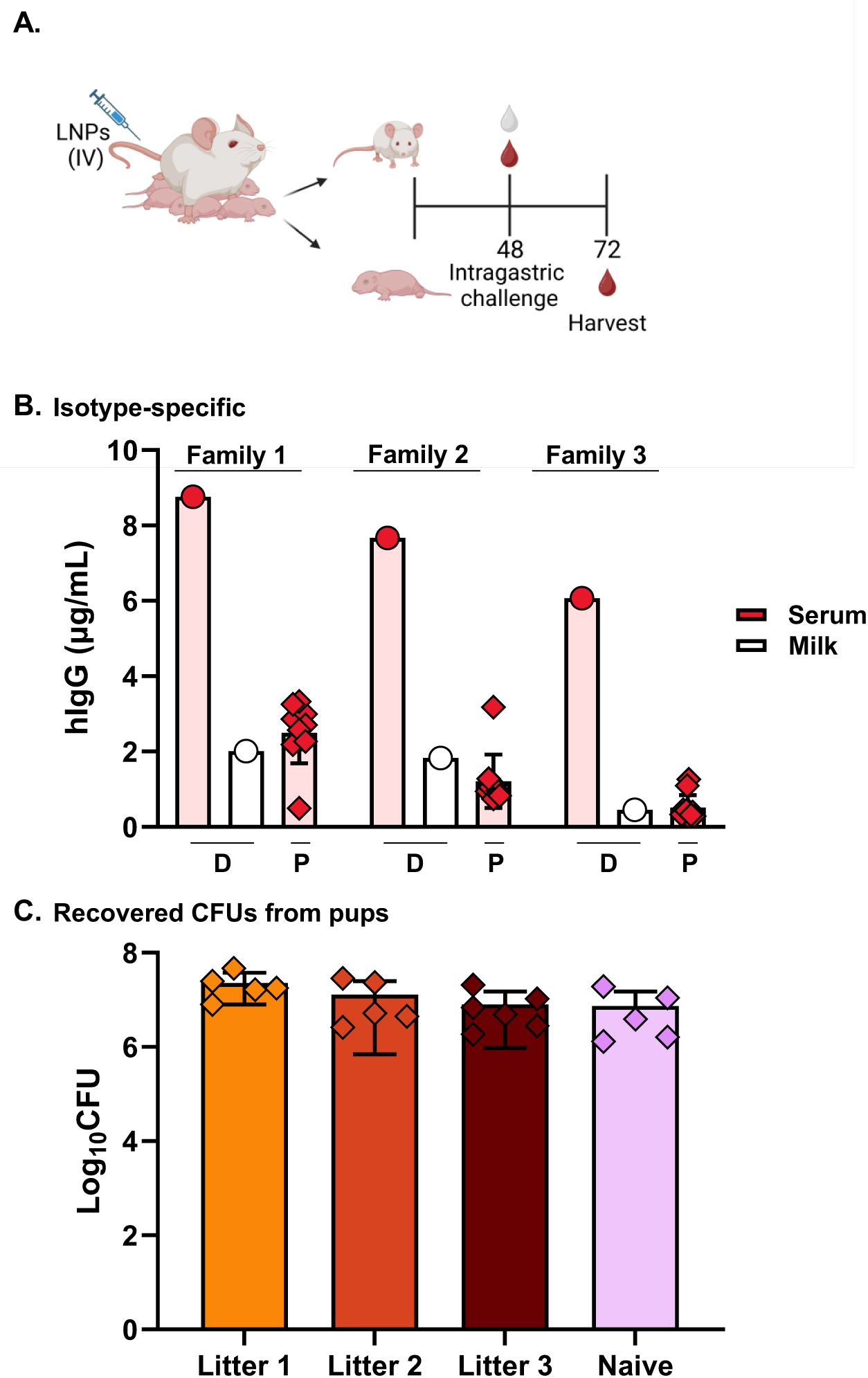
Maternal transfer of ZAC-3 IgG1mRNA to neonates reaches high levels in serum. (**A**) Lactating Swiss Webster mice were intravenously administered with LNPs (1 mg/kg; N = 3) or PBS (N = 2) 3-7 days after giving birth. This was designated as study time 0. 48 h post-administration, pups were intragastrically challenged with *V. cholerae* (1 × 10^5^ CFUs). Serum and milk were collected from dams at 48 h, while serum from pups was collected at 72 h. Pups were euthanized and small and large intestines were harvested, homogenized, and plated for CFUs. (**B**) Concentration of hIgG in Dam serum (pink bars with red circle symbols), milk (white bars with red circle symbols) and pup serum (pink bars with red diamond symbols) were assessed using isotype-specific ELISA. (**C**) Recovered CFUs of *V. cholerae* in the intestinal homogenates from each litter in the ZAC-3 treatment and naïve groups. Dam serum and milk samples are denoted by “D” under the appropriate bars and pup serum samples are denoted by “P”. Bars for dam analysis represent an individual dam while bars for pups show the mean ± SD (n=5-11 mice for the experimental group, n=5 for the naïve group).

## DISCUSSION

The immediate protection afforded by passive immunity has saved millions of infant lives across the globe. For instance, research indicates that SARS-CoV-2 mRNA vaccinations during and after pregnancy conferred significant protection to newborns and reduced hospitalizations through transplacental antibody transfer and/or breastmilk derived neutralizing antibodies [16]. Several studies have shown that at least 4 weeks are required to derive sufficient levels of SARS-CoV-2 neutralizing antibodies in breastmilk because an adaptive immune response had to be generated in the mother following vaccination [17]. However, this may not be rapid enough to protect newborns against infections in emergency situations. The use of mRNA-mediated antibody therapy represents a rapid strategy for the delivery of robust mAb titers to newborns via breastmilk as it eliminates the need for the generation of an adaptive immune response in the mother and overcomes the safety challenges associated with the direct administration of mAbs in the infant. [18]

By employing a previously established mouse model and the well-studied anti-core LPS *V. cholerae* antibody [19], ZAC-3, our study serves as an important proof-of-concept, demonstrating that mRNA-encoded IgG1 mAb can be expressed in a mother and then transferred to her suckling infant via breastmilk. To begin, we confirmed that systemic delivery of mRNA/LNPs enabled the *in-vivo* synthesis and expression of the encoded ZAC-3 IgG in the serum of female mice as early as 1 day after vaccination, with levels sustained for up to 7 days. Additionally, ZAC-3 IgG was detected in breastmilk collected 24 h after mRNA/LNP administration, with concentrations progressively increasing over the course of the study. Furthermore, an equivalent concentration of ZAC-3 in breastmilk was also detected in the serum of breast-fed pups at all time points tested. We also confirmed that ZAC-3 IgG from all biological samples, including the breastmilk and serum of lactating dams and pups, demonstrated vibriocidal activity *in-vitro*.

Our mRNA antibody synthesis and expression system provided sufficient neutralizing antibody titers soon after drug delivery; an important consideration for the implementation of therapeutic antibodies as a treatment platform [20]. To address if the rapid production of antibodies yielded protection in breastfed pups, we analyzed their gut tissues for the extent of colonization by *V. cholerae*. According to our data, no reduction in bacterial burden was observed despite the confirmation of functional activity *in-vitro*. We attribute this observation to the absence of ZAC-3 IgG in the gut tissues of suckling pups where *V. cholerae* primarily colonizes. We hypothesize that the absence of ZAC-3 IgG in the breastmilk-filled guts of the pups was due to the active transport of ZAC-3 IgG from the mucosal tissues in gut to the blood stream. During infant intestinal development, there are two main mechanisms for the systemic absorption of antibodies and other macromolecules. One such method is gut permeability, where protein or drug permeability is size restricted (up to 70 kDa) and depends on the distance of the tissue from the stomach [21]. This form of transport typically occurs before gut closure, a process that occurs around 7 days in human infants and 3 weeks in neonatal mice. The other mechanism involves the active transport of IgG and other immune complexes from mucosal tissues into circulation through binding to FcRn [22]. In rodents, this mechanism is used exclusively to maintain adequate levels of IgG in the blood over extended periods of time [23]. Given the size of IgG and the characteristics of FcRn transport, it is likely that ZAC-3 was actively transported directly into circulation; thus, resulting in undetectable levels of the IgG antibody in the gastrointestinal tract.

IgA unlike IgG is known to provide protection against infection at mucosal surfaces and can be translocated systemically to mucosal sites such as mouse intestines and breastmilk in non-human primates [24]. IgA mAbs also have enhanced antigen binding and neutralization capacity against enteric and respiratory pathogens compared to IgG mAbs of the same clone, making them an excellent candidate for the disease model used in our study. The *in-situ* production of functional, dimeric IgA that can traffic to mucosal sites is possible with the mRNA/LNP technology reported herein. However, evidence from literature shows that systemically delivered dimeric IgA is quickly eliminated due to the short half-life (6-18 h) compared to IgG (15-30 days) [12]. We also found this to be true in our study, as mRNA-encoded ZAC-3 IgA was present at low concentrations in serum and breastmilk and absent in the gut tissues of both pups and dams after systemic delivery (data not shown). This makes working with IgA more challenging because it either needs to be delivered repeatedly and within the window of infection or modified to sustain its presence at the infection site to achieve protection [25]. Despite the lack of *in-vivo* protection by IgG in this specific disease model, the use of an mRNA-encoded IgG has many applications especially for systemic infections in infants including neonatal sepsis, pneumonia and meningitis. Moreover, this study represents the first demonstration, to our knowledge, of maternal transfer of an mRNA-encoded antibody to pups. Furthermore, the expressed IgG maintained binding functionality and neutralization of target pathogen *in-vitro*.

Studies have shown that pregnancy and lactation affect the biodistribution of antibodies and redirect certain antibodies from maternal circulation to the placenta and breastmilk respectively. In light of this, compared to non-lactating mice, we observed a significant drop in the serum ZAC-3 IgG concentration of lactating dams in our study, suggesting an extravascular sink to the mammary glands. Moreover, approximately a fifth of the concentration of ZAC-3 IgG found in serum of lactating dams was detected in the breastmilk. Similarly, a study showed that after SARS-CoV-2 vaccination, IgG antibodies were generally higher in serum than in maternal milk of the same woman. [26]. The biodistribution dynamics of mRNA vaccines and especially mRNA mediated antibody therapies during pregnancy or breastfeeding are poorly investigated [27]. However, evidence in the literature suggests that vaccine mRNA can cross the blood-breastmilk barrier in humans and are eliminated from breastmilk within 3 days of vaccination – albeit at lower concentrations. The mRNA is said to be concentrated in the breastmilk extracellular vesicles with an absence of translational activity [28]. If this were to be the case in our study, it could explain the differences in concentration we see between lactating and non-lactating mice. There’s also a possibility that the expressed IgG in circulation was transported to the breastmilk by active transport via FcRn or by some other means in lactating dams. However, we did not investigate if there’s a direct correlation between the depletion of IgG in the maternal serum and its accumulation in the mammary glands or breastmilk to make this claim. Furthermore, based on the estimated total blood volume in animals, the daily concentration of ZAC-3 in the milk of lactating mice (2-4 µg/mL) did not fully account for the difference in total serum antibody concentration between lactating (∼24 µg total) and non-lactating mice (∼108 µg total). This suggests that other extravascular sites in lactating mice may contribute to antibody accumulation beyond the mammary tissue itself.

The potential applications of an mRNA/LNP-encoded monoclonal antibody technology is significant. Studies have shown the successful use of mRNA antibodies to prevent viral infections e.g. Rabies, Influenza, Zika, HIV-1 and Chikungunya virus [18; 20; 29; 30; 31; 32]. Others have reported success with its use against bacterial infections e.g. *Salmonella*, *Pseudomonas aeruginosa*; botulinum toxin poisoning or toxin related deaths in mice [20; 24]. There are also implications for the use of monoclonal antibodies to precisely modify the human microbiome [33].

In conclusion, this study confirms that to protect against neonatal diseases, the passive immunization of mothers with mRNA/LNP-encoded mAb enables the rapid, safe, sustained and effective delivery of neutralizing monoclonal antibodies to newborns via breastmilk. This unique strategy of passive immunization of newborns further eliminates the safety, pharmacokinetic and clinical pharmacology challenges such as routes of administration, dosing complexities, age and biodistribution often associated with the direct delivery of mAbs to infants. Future studies will focus on the rational design and delivery of mRNA-encoded dimeric IgA or modified IgG through breastmilk to either protect against mucosal associated diseases or reshape the gut microbiota with the overall goal of improving health and survival outcomes in newborns via passive immunization.

## ACKNOWLEDGEMENTS

We thank Dr. Graham Willsey for technical assistance and for valuable feedback and Grace Freeman-Gallant for statistical consultation. We are grateful to the Wadsworth Center’s Veterinary Sciences staff for assistance in oversight of maternal-pup studies. We also thank the Wadsworth Center’s Tissue Culture and Media core facility for bacterial culture media.

---

**Table S1**
| **<u>Group</u>** | **<u>Treatment</u>** | **<u>hIgG (μg/mL), ± SD</u>** |  |  |  | **<u>Vibriocidal Activity (CFUs), ± SD</u>** |  |  |  |
| --- | --- | --- | --- | --- | --- | --- | --- | --- | --- |
|  |  | **<u>24h</u>** | **<u>48h</u>** | **<u>72h</u>** | **<u>168h</u>** | **<u>24h</u>** | **<u>48h</u>** | **<u>72h</u>** | **<u>168h</u>** |
| Naïve | PBS | <LOD | <LOD | <LOD | <LOD | 2.1E+06 | 2.2E+06 | 1.5E+06 | 1.5E+06 |
| Lactating | mRNA | 13.235<br>(2.291) | 15.177<br>(2.876) | 13.719<br>(2.773) | 1.833<br>(0.427) | 260 | <LOD | 1.1E+04 | 1.8E+06 |
| Nonlactating | mRNA | 52.936<br>(13.547) | 62.837<br>(17.752) | 69.535<br>(21.221) | 44.424<br>(11.224) | <LOD | <LOD | <LOD | <LOD |

